# Integrated Genomic Analysis of Mitochondrial RNA Processing in Human Cancers

**DOI:** 10.1101/080820

**Authors:** Youssef Idaghdour, Alan Hodgkinson

## Abstract

Alterations to mitochondrial function and mutations in mitochondrial genes have been reported for a wide variety of cancers, however the mitochondrial transcriptome remains largely unexplored in cancer despite an emerging appreciation of the role that post-transcriptional regulation plays in the etiology of these diseases. Here, we quantify and assess changes to mitochondrial RNA processing in human cancers using integrated genomic analysis of RNA Sequencing and genotyping data from 1226 samples across 12 different cancer types. We find significant changes to m^1^A and m^1^G post-transcriptional methylation rates at functionally important positions in mitochondrial tRNAs in tumor tissues across all cancers. Pathways of RNA processing are strongly associated with methylation rates in normal tissues (P=2.85×10^-27^), yet these associations are lost in tumors. Furthermore, we report 18 gene-by-disease-state interactions where altered methylation rates occur under cancer status conditional on genotype, implicating genes associated with mitochondrial function or cancer (e.g. *CACNA2D2, LMO2* and *FLT3*) and suggesting that nuclear genetic variation can potentially modulate an individual’s ability to maintain unaltered rates of mitochondrial RNA processing under cancer status. Finally, we report a significant association between the magnitude of methylation rate changes in tumors and patient survival outcomes. These results highlight mitochondrial post-transcriptional events as a clinically relevant mechanism and as a theme for the further investigation of cancer processes, biomarkers and therapeutic interventions.

## Introduction

The role of mitochondria in cancer has long been controversial. Although mitochondria are essential for tumor cell growth (1–3), many lines of evidence indicate that altered mitochondrial bioenergetics are required for tumor initiation and persistence. First, the up-regulation of anaerobic energy production via glycolysis, the so-called Warburg effect, is well documented and recognized as a hallmark of cancer (4, 5). Second, mutations in nuclear encoded mitochondrial genes have been identified in patients with cancer, with links to the disease well established in some cases (6). Third, increased numbers of mutations are consistently found in the mitochondrial genomes of tumor cells when compared to normal samples (7–9). These mutations may merely tag carcinogenesis, but whether other genetic properties of mitochondrial genomes are important in tumorigenesis remains one of the important unanswered questions in cancer biology.

In line with this, recent studies have looked beyond mitochondrial DNA mutations to consider other important genetic processes. Mitochondrial copy number has been found to vary between paired normal and tumor samples, with low numbers generally observed in cancer tissue (10); previous work has also highlighted the higher rates of mitochondrial duplication and transcription in tumors (11). Furthermore, Stewart *et al* (12) recently observed mutations in mitochondrial transfer RNA (tRNA) genes with higher alternative allele frequencies in tumor mitochondrial RNA compared to corresponding mitochondrial DNA from the same individuals, suggestive of altered processing of mitochondrial RNA as some transcripts accumulate differently in cancer tissues. However, it is not known whether similar processes also occur in healthy individuals and therefore elucidating the importance of these events in cancer requires further investigation.

Even so, the idea that post-transcriptional processing of the mitochondrial transcriptome may be altered in disease is intriguing. Mitochondrial RNA is transcribed as continuous polycistrons, which are then processed under the ‘punctuation model’, whereby tRNAs that intersperse mRNAs are targeted for modification and cleavage by nuclear encoded proteins (13–16). The polycistronic nature of mitochondrial transcription means that post-transcriptional events are particularly important: knockdown of RNA processing enzymes influences mitochondrial mRNA and protein levels, and mitochondrial function (17) and the rate of m^1^A and m^1^G post-transcriptional methylation at the 9^th^ position of mitochondrial tRNAs (p9 sites) may dictate downstream metabolic phenotypes (18). Indeed, p9 methylation is thought to influence the correct folding of mitochondrial tRNAs, thus influencing the rate of cleavage within the polycistronic transcript and potentially impacting upon their downstream roles in protein translation (19–21). There are several reasons to believe that altered processing of mitochondrial RNA, hereinafter referring to post-transcriptional nucleolytic processing and nucleotide modifications, may be involved in cancer. For example, mutations within the mitochondrial processing enzyme RNase Z were found to be segregating with prostate cancer incidence in human pedigrees (22) and mutations within mitochondrial tRNAs, which are heavily post-transcriptionally modified, have been previously linked with cancer (23). Despite these observations, no large-scale analysis of tumor specific post-transcriptional processing of mitochondria has ever been carried out.

Here, we assess whether mitochondrial RNA processing is altered in cancer by analyzing RNA sequencing data from 1226 paired normal and tumor samples across 12 cancer types from The Cancer Genome Atlas (TCGA). We find significant and consistent signatures of increased mitochondrial tRNA methylation in tumor tissue when compared to paired adjacent normal tissue. These changes are associated with differential mitochondrial RNA cleavage and mitochondrial gene expression profiles, indicative of altered mitochondrial RNA processing in cancer. Furthermore, the changes in mitochondrial tRNA methylation in tumor cells is associated with specific altered nuclear gene expression signatures. We also find evidence of context-specific SNPs that are associated with p9 methylation rates in tumor, but not normal samples (gene-by-disease-state interactions). Finally, we find a significant relationship between the magnitude of change in mitochondrial tRNA methylation in tumor tissue compared to adjacent normal tissue and the survival outcome of patients with Kidney renal clear cell carcinoma, highlighting the potential importance of these events in tumorigenesis.

## Results

### Post-transcriptional changes in tumor cells

To study the patterns of mitochondrial RNA processing in human cancers we mapped and filtered raw RNA sequencing data from 1226 samples from matched tumor-normal pairs across 12 cancer types from The Cancer Genome Atlas (TCGA) (Figure 1A). Using these data we inferred the level of m^1^A and m^1^G post-transcriptional methylation occurring at eleven functionally important positions within mitochondrial tRNAs (the 9^th^ position of eleven different tRNAs, as identified in (18), henceforth referred to as p9 sites) by using the proportion of mismatches observed at these positions, in line with approaches taken by previous studies (17, 18, 24). For each of the eleven p9 sites and within each of the 12 cancers (11×12=132 sites in total), we then compared the level of methylation observed between between paired normal and tumor samples.

**Figure 1:**
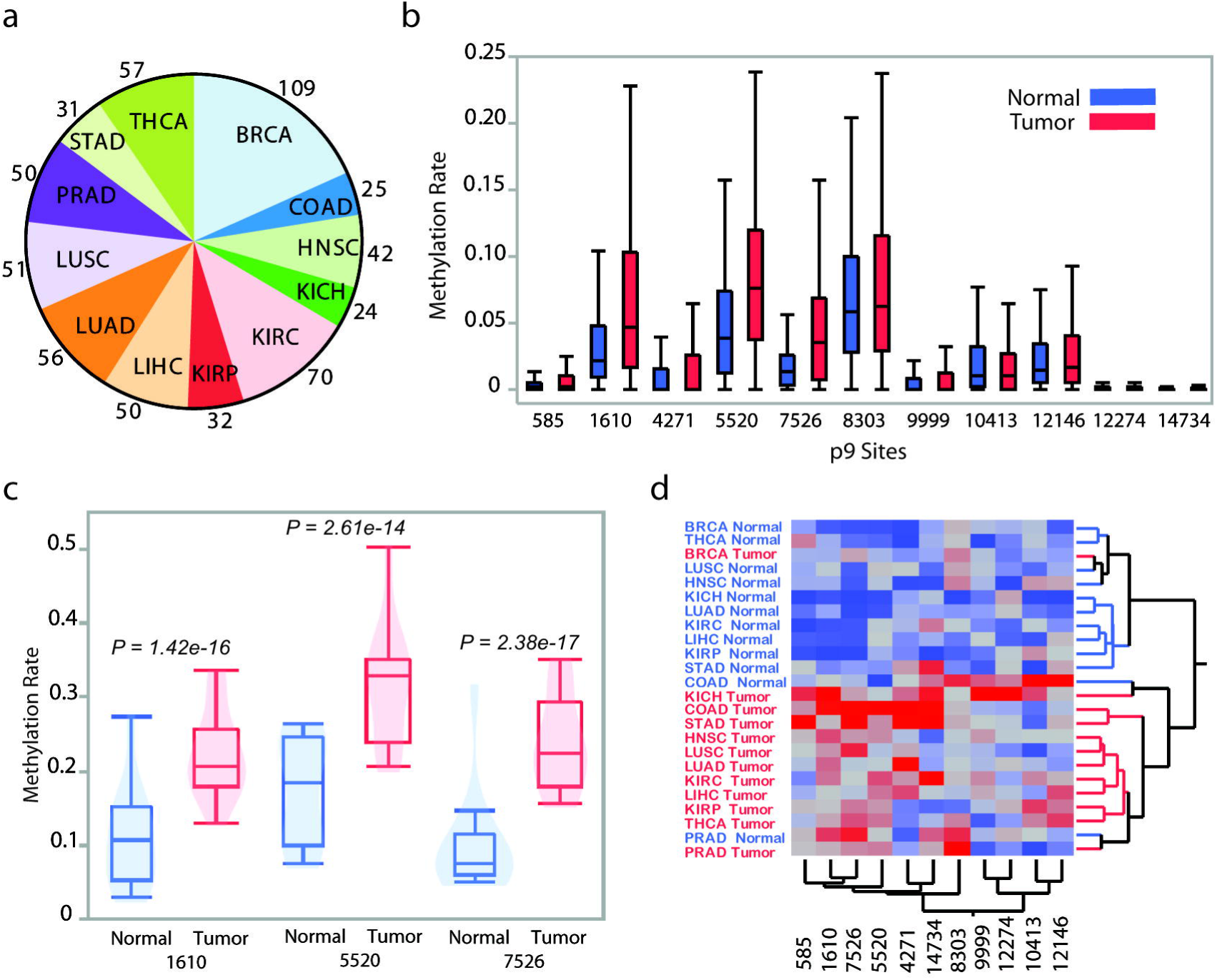
Methylation differences between paired tumor and normal samples at tRNA p9 sites within mitochondrial tRNAs. **(a)** Number of normal-tumor pairs for each cancer type, including Breast invasive carcinoma (BRCA), Colon adenocarcinoma (COAD), Head and Neck squamous cell carcinoma (HNSC), Kidney Chromophobe (KICH), Kidney renal clear cell carcinoma (KIRC), Kidney renal papillary cell carcinoma (KIRP), Liver hepatocellular carcinoma (LIHC), Lung adenocarcinoma (LUAD), Lung squamous cell carcinoma (LUSC), Prostate adenocarcinoma (PRAD), Stomach adenocarcinoma (STAD) and Thyroid carcinoma (THCA), **(b)** Observed methylation rates for all eleven p9 sites within KIRC, split into normal and tumor, **(c)** Standardized methylation rates split into normal and tumor for all cancers combined for p9 sites 1610, 5520 and 7526 and **(d)** Two-way hierarchical clustering of mean standardized methylation rates for normal and tumor samples across all cancer types. The clustering was generated with the Ward method.

In total, 52/132 sites show significant differences between normal and tumor tissues at a 5% significance level (paired t-tests) and 46 of these sites show increases in the observed methylation rate in cancer tissue (Figure 1B for examples showing all eleven p9 sites in KIRC, Supplementary Table 1). After applying Bonferroni correction (within each cancer type, P<0.0045), 21 sites remain significant, 20 of which show increases of methylation rate in cancer tissues. Resampling sequencing reads to the same depth in paired normal and tumor samples (thus accounting for potential biases in sequencing coverage) gives very similar results (Supplementary Table 2, see methods). These observations strongly suggest an increase in the level of post-transcriptional methylation of mitochondrial tRNAs across a large number of sites in tumor tissue from multiple cancer types.

Next, we investigated whether the observed differences in methylation rates are a general trend in cancer. For each p9 site we standardized the data within each cancer (but not across normal or tumor sample types, thus maintaining cancer associated patterns in methylation rates) and tested differences between tumor and normal pairs across all cancer types combined. In total, 8 out of the 11 p9 sites show highly significant differences (P<0.0045, Supplementary Table 3, Figure 1C for p9 sites 1610, 5520 and 7526). Following this, we averaged methylation rates by group and find that, strikingly, 2-way hierarchical clustering using all p9 sites data from 24 cancer-sample type sets revealed how groups cluster largely by sample type (normal or tumor) with the exception of BRCA Tumor, COAD Normal and PRAD Normal not clustering within their respective sample type (Figure 1D). These results suggest that altered processing of mitochondrial RNA is a consistent trend across multiple cancer types.

To infer the impact of changes at sites and cancers where we observe significant differences (at P<0.05), we compared the rates of methylation across both normal and tumor samples with the rate of cleavage occurring at the 5’ end of each respective tRNA, which we measured as the proportion of sequencing reads starting or ending either side of this position. As a control, we considered cleavage rates a further 9bp upstream from each p9 site. Methylation rates significantly correlate with cleavage rates for 10/45 comparisons after Bonferroni correction (P<0.001, vs 1 significant correlation for the control) and for 18/45 comparisons at a 5% significance level, with all significant correlations being in the positive direction (Supplementary Table 4). Furthermore, we considered the influence of changes in tRNA methylation in cancer on mitochondrial gene expression: in normal samples, p9 methylation rates significantly correlate with mitochondrial gene expression in 25 comparisons across cancer types (Supplementary Table 5, Spearman Rank P<0.05 after Bonferroni correction within cancer type, mostly negatively correlated with *MTCO3, MTCO2* and *MTCO1* abundance and positively correlated with *MTND2* abundance), yet in tumor samples these relationships break down and only two pairwise comparisons are significant. In all, these analyses further indicate that major changes to mitochondrial RNA processing take place in human cancers.

### Nuclear transcriptional signatures associated with changes in mitochondrial RNA processing

Mitochondrial RNA transcription and processing, like many other molecular processes taking place in mitochondria, is under strong nuclear control. As such, we tested the hypothesis that the expression of nuclear-encoded genes involved in mitochondrial RNA processing is altered in tumors by performing differential expression analysis. Raw RNA sequencing data were aligned, filtered and normalized as detailed in the Methods and expression data for 99 mitochondrial RNA-binding proteins (as listed in (25)) were retrieved and compared between normal and tumor samples. In this section we report results for BRCA only, since this is the cancer type for which we have the most paired samples (>100) and thus the most power, however we also see the same broad trends in other cancer types with the next largest sample sizes (see methods). In total, we detected 55 genes differentially expressed at Bonferroni significance (Fig 2A) and the heatmap of all 99 gene expression traits surveyed shows that samples cluster largely by sample type (normal or tumor, Supplementary Figure 1).

**Figure 2:**
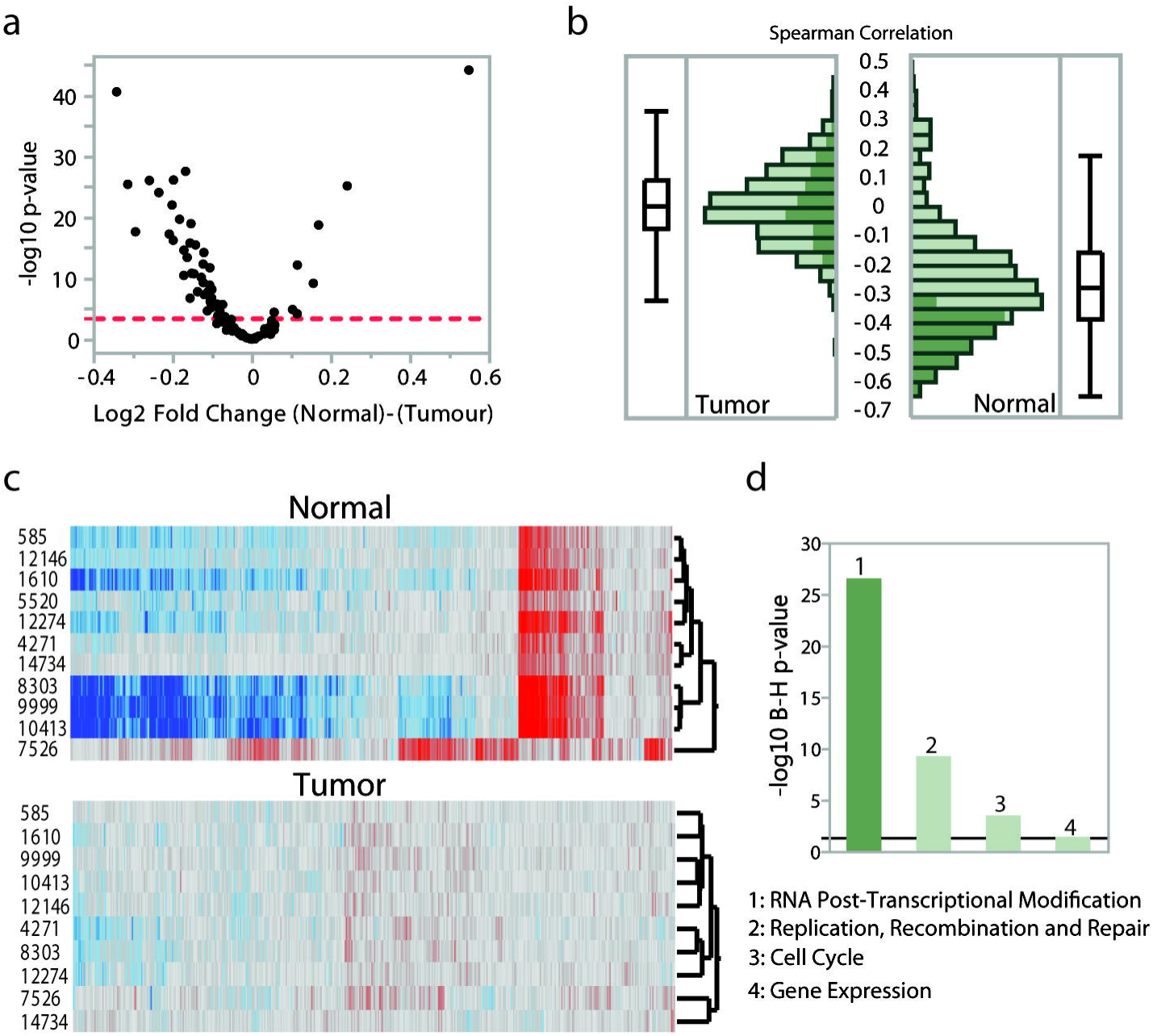
Differential expression and correlations between tRNA p9 methylation rates and nuclear gene expression in BRCA. **(a)** Volcano plot of statistical significance (shown as the negative logarithm of the p-value on the y-axis) versus magnitude of differential gene expression (shown as the log base 2 of magnitude of mean expression difference on the x-axis) of 99 genes encoding mtRNA-binding proteins. The dashed line indicates Bonferroni statistical significance. **(b)** Distribution of Spearman correlations between gene expression levels of the 99 genes and methylation rate at p9 site 10413. Associations that are significant at Bonferroni threshold in Normal samples are highlighted with the dark green color. **(c)** Two-way clustering of Spearman correlations of expression levels of 16,736 genes (columns) and methylation rate at 11 p9 sites (rows) in the BRCA dataset. Correlation values are visualized using a red-to-gray-to-blue color theme (values range from 0.62 to -0.66). **(d)** Ingenuity pathway enrichment analysis of nuclear genes whose expression levels are associated with methylation rate at p9 site 10413 in the BRCA dataset. Biological functions enriched at Benjamin-Hochberg significance threshold are shown.

Following this, we tested for associations between the expression levels of the 99 nuclear-encoded factors and the observed changes in mitochondrial RNA methylation. We performed cross-correlation analysis (Spearman Rank) across all individuals for all possible 1,089 p9 site methylation rate-nuclear gene expression trait pairs in normal and tumor samples separately. The test revealed significant associations for 8 of the 11 p9 sites in normal samples and the total cumulative number of Bonferroni-significant associations was 369 across all eight p9 sites (P<0.0005, Figure 2B for distributions of coefficients, variance explained ranges between -0.64 and 0.47). In sharp contrast, no significant associations were detected in tumor samples (Figure 2B), indicating major deregulation of these processes in cancer. A 2-way hierarchical clustering heatmap of the full correlation matrix (Supplementary Figure 2) shows the consistency of the associations across different p9 sites in normal samples, whereas these associations are highly perturbed under cancer status.

Next, we identified cell-wide processes associated with the observed changes in mitochondrial RNA processing by performing global cross-correlation analysis between the expression levels of all nuclear genes and the methylation rate at each p9 position using the same strategy as outlined above and using a Bonferroni corrected p-value threshold of 3×10^−6^ (00.5/16,736). In normal samples, the test revealed an average of 2311 significant associations for 8 of the 11 p9 sites (Figure 2C, Supplementary Table 6, three sites show no significant associations, variance explained ranges between -0.66 and 0.62). To investigate the functional characteristics of nuclear genes whose expression levels are associated with methylation rate at the p9 site showing the strongest signal in normal cells (p9 site 10413, 6061 genes) we used Ingenuity Pathway Analysis. We find that “RNA Post-Transcriptional Modification” is the top and most highly enriched molecular and cellular function category with 5 significantly enriched sub-functions (Benjamini-Hochberg P-value range 8.48×10^−6^ – 2.85×10^−27^, Figure 2D, “Processing of RNA” was the most enriched sub-function), supporting the idea that many nuclear genes play a role in mitochondrial RNA processing. However, we again find striking differences in tumor samples, where we detect only 2 significant associations across all p9 sites, pointing to major deregulation of nuclear-associated mitochondrial RNA processing in cancer (Supplementary Figure 3).

**Figure 3:**
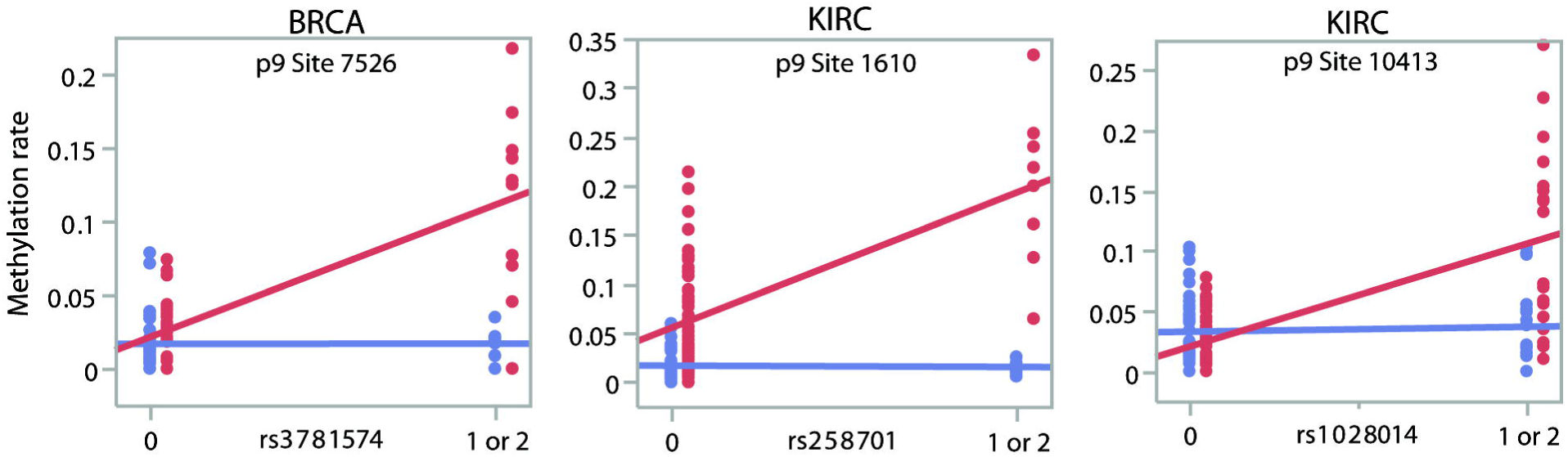
Interaction effects on tRNA p9 sites methylation. The three plots show examples of SNPs that have different relationships with p9 methylation rates in normal (blue) and tumor (red) samples.

### Joint action of genotype and cancer state on post-transcriptional methylation

Given the general increase in p9 methylation in cancers, we assessed whether nuclear genetic variants could modulate the observed changes in mitochondrial RNA processing differentially in tumor relative to normal samples. To do this, we looked for genotype-by-state (tumor or normal) interaction effects on p9 methylation rates across cancer types. We obtained genotyping data for the same samples for which we inferred p9 methylation rates, and limited the analysis to SNPs in Hardy-Weinburg Equilibrium (P>0.001), those where at least 40 individuals had methylation data available and with at least five individuals carrying a minor allele. Under a dominant model and after filtering SNPs for MAF>5%, this analysis identified 18 peak genotype-by-state interactions at genome-wide significance across cancer types and p9 sites (Table 1, and Figure 3 showing examples). In 15 cases the minor allele is associated with increased methylation rates in tumor but not normal samples, suggesting that mitochondrial RNA processing is affected differently in individuals carrying these alleles under cancer status.

**Table 1:** Interaction effects on tRNA p9 sites methylation rate

| Cancer | p9 Site | rs Number | Chr | Position | Location | Gene | N | MAF | P-value |
| --- | --- | --- | --- | --- | --- | --- | --- | --- | --- |
| BRCA | 7526 | rs3781574 | 11 | 33885268 | intron | LMO2 | 46 | 0.161 | 4.61E-11 |
| BRCA | 7526 | rs17690328 | 11 | 33885390 | intron | LMO2 | 46 | 0.135 | 1.17E-08 |
| BRCA | 7526 | rs317391 | 17 | 32174605 | intron | ASIC2 | 44 | 0.085 | 2.46E-09 |
| KIRC | 1610 | rs258701 | 7 | 81761122 | intron | CACNA2D1 | 61 | 0.058 | 8.27E-10 |
| KIRC | 8303 | rs10266772 | 7 | 14132727 | Intergenic | --- | 61 | 0.080 | 2.27E-08 |
| KIRC | 10413 | rs1028014 | 8 | 40698232 | intron | ZMAT4 | 64 | 0.174 | 2.88E-08 |
| KIRC | 12146 | rs17164416 | 5 | 99387447 | Intergenic | --- | 65 | 0.051 | 9.95E-09 |
| LIHC | 12146 | rs41465346 | 1 | 219059708 | Intergenic | --- | 48 | 0.052 | 3.73E-08 |
| LIHC | 12146 | rs16926871 | 8 | 62027498 | intron | CLVS1 | 47 | 0.074 | 1.24E-08 |
| LUAD | 585 | rs13029285 | 2 | 205916078 | intron | PARD3B | 49 | 0.057 | 1.81E-10 |
| LUAD | 10413 | rs7328699 | 13 | 28615701 | intron | FLT3 | 46 | 0.107 | 9.15E-10 |
| LUAD | 10413 | rs12591927 | 15 | 31833911 | intron | OTUD7A | 44 | 0.077 | 2.44E-08 |
| LUAD | 10413 | rs17003208 | 22 | 43073031 | Intergenic | --- | 45 | 0.074 | 1.13E-08 |
| LUAD | 10413 | rs17003212 | 22 | 43079754 | Intergenic | --- | 46 | 0.063 | 1.18E-11 |
| LUAD | 12146 | rs4741498 | 9 | 15424938 | intron | SNAPC3 | 45 | 0.063 | 3.68E-08 |
| LUAD | 12146 | rs7026970 | 9 | 15445219 | intron | SNAPC3 | 45 | 0.063 | 3.68E-08 |
| LUAD | 12146 | rs7046713 | 9 | 15458921 | intron | SNAPC3 | 45 | 0.063 | 3.68E-08 |
| LUAD | 14734 | rs341737 | 8 | 2782370 | Intergenic | --- | 54 | 0.107 | 9.86E-09 |
| PRAD | 10413 | rs7631369 | 3 | 1028679 | Intergenic | --- | 48 | 0.070 | 3.45E-08 |
| PRAD | 10413 | rs7301597 | 12 | 30175147 | Intergenic | --- | 48 | 0.050 | 3.85E-09 |
| THCA | 585 | rs17033484 | 4 | 156686737 | intron | GUCY1B3 | 54 | 0.089 | 3.78E-08 |
| THCA | 4271 | rs2145836 | 20 | 47421064 | intron | PREX1 | 44 | 0.054 | 4.41E-08 |

After initial discovery, we attempted to replicate each genome-wide significant interaction effect in other cancer types at any p9 site (P<0.001, under the same criteria outlined above). In doing so, we find an interaction effect for rs317391 in KIRC at p9 site 12146(P=2.16×10^−5^, interaction effect originally observed in BRCA at p9 site 7526, P= 2.46 ×10^−9^). This SNP falls within an intron of the gene ASIC2, which is part of a sodium channel superfamily. We also find that rs341737 is an interaction SNP for methylation rates at p9 site 14734 in PRAD (P=0.000745, interaction effect original observed in LUAD at p9 site 14734, P=9.86 ×10^−9^). This SNP falls in an intergenic region near to CMSD1, a known tumor suppressor gene. The fact that these SNPs are also associated with similar effects in other cancer types suggests that our initial observations are robust.

### Biological implications of altered post-transcriptional events

In order to relate changes in mitochondrial RNA processing to potential biological outcomes we used cox proportional hazards tests to determine if changes in p9 methylation rates between paired normal and tumor samples are a significant predictor of patient survival outcomes. To ensure power to detect significant associations, we focused on cancers where we had data for at least 50 individuals with >25% death rate within 60 months of diagnosis (KIRC and LUAD). Treating p9 methylation differences as a quantitative trait we find that methylation differences do not significantly predict patient survival in LUAD. However for KIRC, seven p9 sites are significant predictors of patient survival (P<0.05, Supplementary Table 7), with larger increases in p9 methylation rates in tumor compared to paired normal samples being associated with worse survival. The strongest effect occurs for p9 site 10413 (9^th^ position of TRNR, P=0.000476). Treating the change in methylation rate as a categorical variable with data binned into two equal sized groups of high and low methylation differences (Figure 4, P=0.036), the model suggests that those in the larger methylation differences group are 2.784 (95% confidence interval (1.07,7.25)) times more likely to die over a 60 month period after diagnosis.

**Figure 4:**
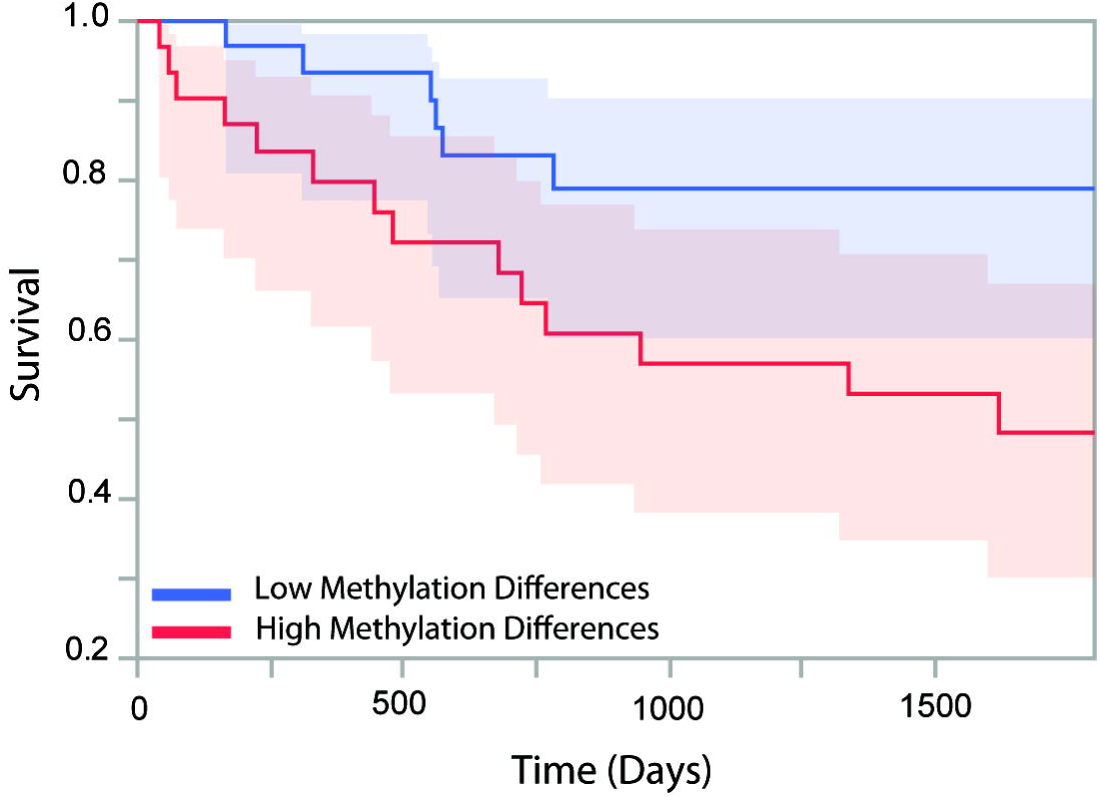
Survival analysis. Data shows relationship between methylation rate differences between paired tumor and normal samples at p9 site 10413 and survival rates for patients with KIRC.

## Discussion

By analyzing RNA sequencing data across a large number of individuals and sample types, we find significant and consistent changes in the rate of m^1^A and m^1^G post-transcriptional modification at functionally important positions within mitochondrial tRNAs in tumor samples when compared to paired normal data. These changes appear to be a widespread phenomenon across different types of tumor and suggest that altered mitochondrial RNA processing is a hallmark of cancer. We hypothesize that oncogenic signals trigger these changes, which in turn serve as a common mechanism by which oncogenetic pathways promote tumorigenesis.

Pathway enrichment analysis of genes implicated in associations between mitochondrial RNA processing and nuclear gene expression unambiguously links methylation rates in mitochondria from normal samples to cell-wide processes of RNA modification and processing. In tumour samples, these relationships completely break down, and focused analysis of genes encoding mitochondrial RNA-binding proteins (25), many of which directly modulate mitochondrial RNA processing, indicates that their misexpression in tumors is driving the observed changes. Proving causality of such events is challenging but is a hypothesis worth pursuing. Based on the fact that mitochondrial tRNAs are key elements in protein translation of genes of the oxidative phosphorylation system (16) and given the connection between cell growth and protein synthesis, it stands to reason that increased rates of tRNA processing would augment the translational and metabolic capacity of mitochondria, modulate the cell cycle and ultimately promote uncontrolled growth. Further studies are warranted to illuminate the molecular mechanisms likely coupling altered levels of mitochondrial RNA processing to cell growth and other tumorigenesis steps such as cell invasion and migration.

The observed differences in the rate of methylation between normal and tumor samples are significant. However, a subset of tumor samples exhibit methylation rates in the range detected in normal samples. As such, we hypothesized that nuclear genetic variation could potentially modulate an individuals’ ability to maintain normal levels of mitochondrial RNA processing. Our test to uncover genotype-by-state interactions revealed 18 peak variants at genome-wide significance that at least partially explain the observed differences between normal and tumor samples. In the vast majority of cases, individuals carrying a minor allele at specific sites saw increased p9 methylation rates in tumor tissues compared to normal, whereas those only carrying the major allele saw no such differences. These in vivo interaction effects serve as a promising starting point for a potential method to uncover allelic variation that conditions the change in mitochondrial RNA processing. Although, no single interaction should be taken as strong evidence of a role for the given SNP in tumor development, among the genome-wide significant interaction SNPs that fall within intronic or exonic regions (11 in total), there are several noteworthy examples where these genes may be interesting for further study. Four genes have a link to mitochondrial function, cellular proliferation or apoptosis (*CACNA2D2, CLVS1, OTUD7A and FLT3*), three genes have been linked to neurological function (and Amyotrophic Lateral Sclerosis in particular), where dysfunctional mitochondria have a known role (26, 27) (*ASIC2, CLVS1, PARD3B*) and four genes have been previously linked with cancer (*LMO2, CACNA2D2, FLT3* and *PREX1*).

Finally, we find that altered mitochondrial tRNA methylation profiles in cancer samples correlate with patient survival outcome in one cancer type. Although these analyses are naturally underpowered to draw firm conclusions about the underlying processes, these results are suggestive of important downstream consequences of altered mitochondrial RNA processing. An alternative hypothesis is that other confounding factors strongly linked to survival may influence mitochondrial RNA processing. Further analysis with larger sample sizes is required to truly understand these relationships.

Taken as a whole, these results highlight the alteration of the mitochondrial post-transcriptional modification and processing landscape taking place in cancer. Furthermore, these observations complement an emerging appreciation of the role that post-transcriptional and post-translational regulation plays in the etiology of cancer (28–31). Although the mechanisms underlying these alterations remain to be resolved, it is tempting to speculate that restoring normal levels of modification and processing of RNA would represent a promising area of investigation for the development of new anticancer therapeutic interventions.

## Materials and Methods

### RNA sequencing data

Institutional approvals from NIH, KCL and NYUAD were obtained to access TCGA data. Raw sequencing files (fastq format) were obtained from TCGA through the CGHub repository via dbGaP accession number phs000178.v9.p8 (32, 33) for twelve cancer types where at least 25 paired tumor and adjacent normal samples were available. These included Breast invasive carcinoma (BRCA), Colon adenocarcinoma (COAD), Head and Neck squamous cell carcinoma (HNSC), Kidney Chromophobe (KICH), Kidney renal clear cell carcinoma (KIRC), Kidney renal papillary cell carcinoma (KIRP), Liver hepatocellular carcinoma (LIHC), Lung adenocarcinoma (LUAD), Lung squamous cell carcinoma (LUSC), Prostate adenocarcinoma (PRAD), Stomach adenocarcinoma (STAD) and Thyroid carcinoma (THCA). In total, we obtained 1226 RNA sequencing datasets for analysis (Supplementary Table 8).

Sequencing reads were trimmed for adaptor sequences, terminal bases with quality lower than 20 and poly-A tails of 5 nucleotides or greater before being aligned to a reference genome (1000G GRCh37 Reference) with STAR 2.51a (34), using default parameters, two-pass mapping and version 19 of the Gencode gene annotation. Careful attention was paid to minimize the likelihood of incorrectly placed reads, particularly those associated with NUMT sequences. To achieve this, a stringent filtering pipeline was applied, as we previously demonstrated (18), focusing only on properly paired and uniquely mapped reads.

### Gene expression levels

To calculate transcript abundances, we used HTseq (35) with default parameters, the ‘intersection-nonempty’ model and Gencode gene annotation file v19. Raw counts were then converted to transcripts per million (TPM). Within the TPM calculation, for mitochondrial genes the total number of fragments mapping to the mitochondrial transcriptome was used to normalize for library size, thus accounting for differences in mitochondrial copy number across samples. For nuclear genes, the total library size was used. TPM scores were then log_10_ transformed and median normalized. Principal component analysis and distribution analysis were used identify outlier samples. Samples greater than three standard deviations from the mean in any of the first three principal components were deemed outliers. All samples paired with these outliers were also removed from subsequent analysis resulting in a set of 1196 samples across all cancers. Distribution analysis shows that samples within each cancer type had similar distributions suggesting that variations due to technical reasons in the data are minimal.

### Methylation rates at tRNA p9 sites

Previous studies have highlighted that sequencing mismatches observed at particular positions in the mitochondrial transcriptome represent post-transcriptional modification events (17, 18, 24). In particular, Hodgkinson *et al* (18) found that the proportion of mismatches at the 9^th^ position of eleven different mitochondrial tRNAs (p9 sites) represents the level of post-transcriptional methylation. Under this assumption, within each sample we inferred the level of p9 methylation by using samtools v1.2 mpileup (36) with default parameters to generate allele count files, from which we calculated the proportion of non-reference alleles at each p9 site (positions 585, 1610, 4271, 5520, 7526, 8303, 9999, 10413, 12174, 12246 and 14734 in the mitochondrial transcriptome). It is important to note that these p9 positions in the mitochondrial transcriptome have previously been shown not to overlap with known variants in NUMT sequences in the human reference using a careful and stringent mapping and filtering strategy (18).

We compared levels of methylation at each p9 site and within each cancer using paired t-tests (132 tests in total) for individuals and sites where the p9 position had at least 20X coverage for both the tumor and paired normal sample. In order to control for any biases in coverage, we repeated the analysis after resampling sequencing reads within each individual and at each site to the lowest coverage found at that site in either the normal or tumor sample. To directly compare p9 methylation across cancers, we standardized rates at each p9 site within each cancer by dividing by the maximum value observed across normal and tumor samples. To compare p9 methylation with cleavage rates at the 5’ end of mitochondrial tRNAs, we calculated the proportion of reads that started or ended either side of the position 9bp upstream of the p9 site compared to all reads covering that position. We considered only sites with at least 20 individuals with 20X coverage at both cleavage and p9 positions and used Spearman rank correlation tests. For comparisons to mitochondrial gene expression, we performed spearman rank correlation tests for each p9 site and within each cancer for normal and tumor samples separately.

### Differential expression and cross-correlations with nuclear gene expression

We evaluated the magnitude and significance of differential expression of transcripts of nuclear-encoded mitochondrial RNA-binding proteins (25) using analysis of variance and Bonferroni thresholds to infer statistical significance. In total, 99 transcripts out of 107 listed in Wolf and Mootha (25) were deemed expressed in BRCA and KIRC datasets (100 transcripts in THCA). Two-way clustering of gene expression data of the full set of genes encoding mitochondrial RNA-binding proteins was generated using Ward’s method in JMP Genomics 8.0 (SAS Institute). To investigate the relationships between nuclear gene expression traits and p9 methylation rate we performed an unbiased genome-wide association between methylation rate at eleven p9 sites and 16,736 expression traits in the BRCA dataset. We calculated Spearman correlation across all individuals in each sample type. The significance of correlations was assessed by correcting for multiple testing resulting in a Bonferroni threshold of 3 × 10^−6^. In the main text we present results for BRCA, however we see the same general trends for THCA and KIRC as significant associations in normal tissue are altered in tumor samples (Supplementary Figure 4). Gene set enrichment analysis was performed using the Core Analysis Workflow implemented in the Ingenuity Pathway Analysis package to measure the likelihood that the association between nuclear genes whose expression was significantly associated with p9 methylation rate and a given process or pathway is due to random chance. The P-value is calculated using the right-tailed Fisher Exact Test that takes into consideration the number of focus genes that participate in that process in question and the total number of genes that are known to be associated with that process in the human reference set. P-values are corrected for multiple testing based on the Benjamini-Hochberg method. The same analysis parameters were used for BRCA, KIRC and THCA (Supplementary Table 9).

### Interaction effects

Where available, we downloaded birdseed genotype files generated from Affymetrix Genome-wide Human SNP arrays (6.0) for all individuals for which we had RNA sequencing data. Samples that did not pass TCGA quality control were not used and in total, data were available for 569 individuals (Supplementary Table 8). For each cancer type, we filtered genotypes with birdseed quality scores above 0.1 and kept SNPs in Hardy-Weinberg Equilibrium (P>0.001). Since sample sizes within cancers were generally small, we converted minor homozygote alleles to heterozygotes (dominant model). SNPs with MAF<5% were removed and we then ran a quantitative trait model (GxE) in plink 1.07 (37), using the rates of p9 methylation at sites where paired normal and tumor samples had at least 20X coverage. Within these tests, p9 methylation rates were used as the quantitative trait, sample type (normal or tumor) as the environment and regression coefficients were compared between normal and tumor association tests to generate a P-value for the interaction term. In order to ensure robust findings, we considered only sites that had data for at least 40 individuals. QQ plots for p9 sites and cancers showing significant interaction effects are shown in Supplementary Figure 5. After identifying SNPs that passed genome-wide significance, we visually inspected data plots and made sure that the uncovered associations are not driven by outliers. The GxE test using paired datasets is robust as both contrasted groups (Normal and Tumor) have the same phenotypes. However, to ensure that age and sex are not affecting the results, we tested whether age or gender are correlated with methylation rates within either normal or tumor samples for individuals that we had genotyping data for; in all cases we find no significant relationships (P>0.05 after Bonferroni correction). SNP annotations were taken from the Affymetrix annotation file associated with the array.

### Survival analysis

We obtained patient survival data from TCGA and performed survival analysis in R using the package ‘Survival’. We limited analysis to cancers for which we had RNA sequencing data for at least 50 individuals, with a death rate of 25% (Kidney renal clear cell carcinoma and Lung adenocarcinoma). Censoring was limited to 60 months, since most events happen during this time. Cox proportional hazards tests were used to model survival as a function of changes in p9 methylation rates in tumor versus normal samples. For significant associations, we tested Schoenfeld residuals to ensure that the proportional hazards assumption was being met (P>0.05 in all cases). To calculate meaningful hazard ratios, tests were repeated after binning changes in p9 methylation rates into two equal sized groups. Results reported in the main text include no covariates. Repeating the analysis including age, sex and ethnicity gives similar results: for KIRC, 5 out of 7 remain significant at P<0.05, and the strongest effect for the 9^th^ position of *TRNR* has a P-value of 0.00151.

## Acknowledgements

The results published here are in whole or part based upon data generated by The Cancer Genome Atlas managed by the NCI and NHGRI. Information about TCGA can be found at http://cancergenome.nih.gov. We thank Seth Seegobin for advice regarding the survival analysis. Y.I. is funded by a New York University Abu Dhabi research grant (AD105). A.H. holds an MRC eMedLab Medical Bioinformatics Career Development Fellowship, funded from award MR/L016311/1. A.H. also holds a WHRI-Academy Marie Curie (COFUND) Fellowship and the research leading to these results has received funding from the People Programme (Marie Curie Actions) of the European Union’s Seventh Framework Programme (FP7/2007-2013) under REA grant agreement n° 608765. Work presented here reflects only the author’s views and not the views of the European Commission. The research was supported by the National Institute for Health Research (NIHR) Biomedical Research Centre based at Guy’s and St Thomas' NHS Foundation Trust and King’s College London. The views expressed are those of the authors and not necessarily those of the NHS, the NIHR or the Department of Health.

The authors declare that they have no competing interests.

